# The Effect of Polymer Length in Phase Separation

**DOI:** 10.1101/2022.11.21.517354

**Authors:** Gilberto Valdes-Garcia, Kasun Gamage, Casey Smith, Karina Martirosova, Michael Feig, Lisa J. Lapidus

## Abstract

Understanding the thermodynamics that drives liquid-liquid phase separation (LLPS) is quite important given the many numbers of diverse biomolecular systems undergoing this phenomenon. Regardless of the diversity, the processes underlying the formation of condensates exhibit physical similarities. Many studies have focused on condensates of long polymers, but very few systems of short polymer condensates have been observed and yet studied. Here we study a short polymer system of various lengths of poly-Adenine RNA and peptide formed by the RGRGG sequence repeats to understand the underlying thermodynamics of LLPS. We carried out MD simulations using the recently developed COCOMO coarse-grained (CG) model which revealed the possibility of condensates for lengths as short as 5-10 residues, which was then confirmed by experiment, making this one of the smallest LLPS systems yet observed. Condensation depends on polymer length and concentration, and phase boundaries were identified. A free energy model was also developed. Results show that the length dependent condensation is driven solely by entropy of confinement and identifies a negative free energy (*-ΔG*) of phase separation, indicating the stability of the condensates. The simplicity of this system will provide the basis for understanding more biologically realistic systems.

## INTRODUCTION

Liquid-liquid phase separation (LLPS) is an increasingly predominant phenomenon found in many types of cells and under many conditions^1-2^. Despite the diversity of the formation conditions and their contents, there seem to be commonalities of resulting condensates that can be described by physics-based models^3^. While condensates have been observed for folded proteins and RNAs^4^, the majority of systems studied are of disordered proteins and nucleic acids. Polymeric models describing intra- and intermolecular interactions balanced against entropy have been developed to describe such observations. The simplest model was proposed by Flory and Huggins to describe the free energy change of mixing a homopolymer with a solvent^5-6^. Pappu and others have expanded this theory into the stickers-and-spacers model to include specific interactions in heteropolymers, while still accounting for the entropy of polymer.^7^ Similarly, Rosen and others have described condensation in terms of valency of client binding to scaffold molecules^8^.

Experimentally, most model systems use proteins and/or RNAs that are dozens to hundreds of residues long^9-13^. Alshareedah, et al.^14^ worked with sequences similar to those used here, but at lengths of 500 bases and 50 amino acids. Bai et al.^10^ demonstrated condensation with a 21-base oligonucleotide and a 30-residue peptide in the presence of inert crowders. Lim et al.^15^ observed condensates in 23-residue peptides rich in histidine and tyrosine. Akahoshi et al.^16^ observed condensation with polyA^15^ ssDNA and K_7_X_3_ peptides, where X represents several different residues. Tang et al.^17^ conducted an exhaustive computational survey of all dipeptide sequences and experimentally demonstrated liquid droplets of QW. Finally, recent work by Fisher and Elbaum-Garfinkle^18^ has shown condensation with UDP or UTP and polyR_10_, but not with UDP and polyK_10_, indicating that arginine has stronger interactions with RNA than lysine, despite the same charge. These differences have been described computationally as well using all-atom simulations for polyR_5_/polyK_5_ – polyUracyl_5_ mixtures at various concentrations^19^. However, the length dependence at the residue level beyond this precision was not explored. Therefore, a natural question is, how long must such polymers be to observe condensation?

We recently developed a coarse-grained (CG) computational model that reduces each amino acid or base to a single bead^20^. The interactions between beads were systematically parameterized to match experimental observations of both polymer characteristics and LLPS for many different systems. The model was also developed to accurately account for concentration dependence of LLPS, which allowed quantitative prediction of the length dependence under real experimental conditions. The accuracy of this model compared to others in the literature was achieved with just a few specific parameters, the stiffness of the angular harmonic potential and separate parameters for the strength of cation-*π* interactions within a protein and between protein and nucleic acids.

Using this model, we predicted that short polymers would undergo LLPS at moderate (∼1 mg/mL) concentrations. Subsequently, we systematically investigated LLPS of RNAs and peptides of various lengths computationally and experimentally. The composition and the volume fraction of the polymers were kept constant so that the only variable that changed was the number of covalent bonds in the system. Here we experimentally observed condensation of molecules as short as 5-10 nucleotides or amino acids, making this one of the smallest LLPS systems yet observed.

## RESULTS AND DISCUSSION

Using COCOMO, our recently-developed CG model^20^, simulations were performed for various lengths of adenine polymers (polyA_N_) and different repetitions, M, of the [RGRGG]_M_ peptide for a fixed volume fraction of 0.13% of each polymer (**Table S2**). Snapshots shown at the end of the simulations indicate system-dependent condensation (**Fig. S1**). According to these initial simulations there are minimum peptide and nucleic acid polymer lengths before condensation is observed. The minimum peptide length required for phase separation depended on the length of polyA_N_ and *vice versa*, the minimum nucleic acid lengths depended on the peptide length. For example, [RGRGG]_4_ was the minimum length to form clusters with polyA_20_ whereas polyA_10_ required at least the length of [RGRGG]_10_ to form clusters (**see Fig. S1**). Moreover, we found that even for RNA as long as 300 bases, no clusters were observed for [RGRGG]_1_ or [RGRGG]_2_, indicating that there is a minimal peptide length required for phase separation. No condensates were observed for polyA_5_ for peptides as long as [RGRGG]_15_, also suggesting a minimal RNA length, but the peptide length was not extended beyond 75 amino acids. These simulations are generally in agreement with previous computational works on RNA^21-23^ and DNA^24^ showing LLPS is enhanced with longer nucleic acids lengths.

To confirm the predictions from the CG model, LLPS was tested for different mixes of polyA_N_ (N = 5, 10 and 20) with [RGRGG]_M_ (M = 1,2,3,4,6,8 and 10). For all experiments the total concentrations of RNA and peptide were maintained at 1 mg/mL. Phase separation was observed either by fluorescence imaging, using Cy3-labeled polyA_N_ or Cy5-labeled [RGRGG]_1_, or by Differential Interference Contrast (DIC) imaging. We observed that the Cy3 in the labeled RNA induces phase separation by itself, likely due to dye hydrophobicity and stacking interactions in oligomers^25^. To find out the threshold at which Cy3 starts inducing phase separation in our systems, the concentration of polyA_10_-Cy3 was varied 0-100 μM in a mixture of polyA_10_ and [RGRGG]_2_. Results showed that no condensates formed up to 20 μM, but condensates were observed at 30 μM and above (**Fig. S2**), indicating that the threshold lies in between 20 and 30 μM. Therefore, Cy3-labeled RNA was kept at a very low concentration of 5 μM compared to unlabeled polyA to prevent it from inducing phase separation.

**Fig. 1** shows the confocal microscopy images for all combinations of [RGRGG]_1,2,3,4,6,8,10_ and polyA_5,10,20_ and long chain polyA (polyA_>600_). Surprisingly, the minimum peptide length for polyA_20_ to phase separate was [RGRGG]_2_, and polyA_10_ separated with [RGRGG]_3_, shorter than predicted from the initial simulations. Additionally, polyA_5_ phase separated with [RGRGG]_4_, whereas the model did not predict phase separation at any peptide length. For each polymer, increasing the length by no more than 5 residues was sufficient to induce phase separation. It was observed by naked eye that all samples with condensation turned cloudy once the peptide was mixed with RNA, but [RGRGG]_4_ and [RGRGG]_6_ with PolyA_5_ were less cloudy compared to all the other mixtures, suggesting that condensation for these mixtures is less stable than for longer lengths. The results also show that no phase separation occurred for the shortest peptide [RGRGG]_1_ even with polyA_>600_, indicating that [RGRGG]_1_ does not interact sufficiently favorably with the RNA polymer to form condensates.

**Figure 1.**
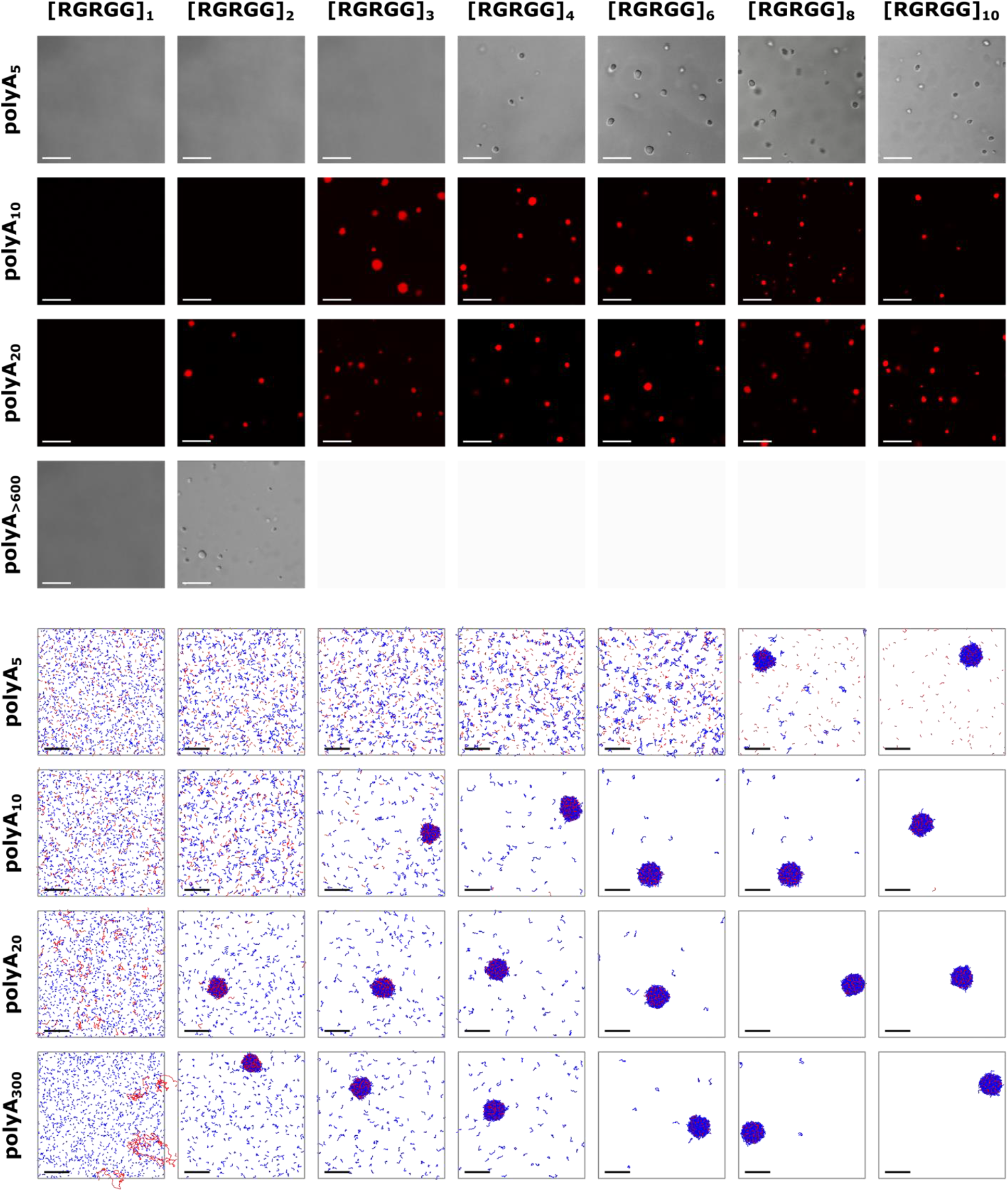
Condensates of various lengths of RNA and peptide. Experimental (upper panels) and simulation using COCOMO 1.2σ (lower panels) results are shown for different mixtures. RNA and peptides at 1 mg/mL. Images were obtained by confocal and DIC microscopy. 5 μM Cy3-labeled polyA was used for fluorescence. The final frames of the simulation trajectories show the central box of the periodic system for each simulated system with RNA and protein were colored in red and blue, respectively. Scale bars represent 10 μm in experimental panels and 20 nm in the simulation panels.

The images in **Fig. 1** shows that there were no substantial differences in droplet size or abundance right after mixing for any of the combinations except for [RGRGG]_4_ and [RGRGG]_6_ with PolyA_5_. To confirm, confocal images were analyzed to determine the area of each condensate. **Fig. 2** shows the histograms of various combinations of RNA and peptide, which were fit to a Poisson distribution. Within the uncertainty of such distribution, there were no substantial differences at the earliest time between different combinations of RNA and peptide. The average condensate size slightly increases with time, but differences in growth do not appear to depend on the lengths of the components. This suggests that the composition of the condensed phase, especially the residue densities, does not change much with polymer length, as discussed below. Nucleation appeared to occur within the experimental time of mixing the components and creating the first confocal image, but it is possible that the rate of nucleation does depend on peptide or RNA length.

**Figure 2.**
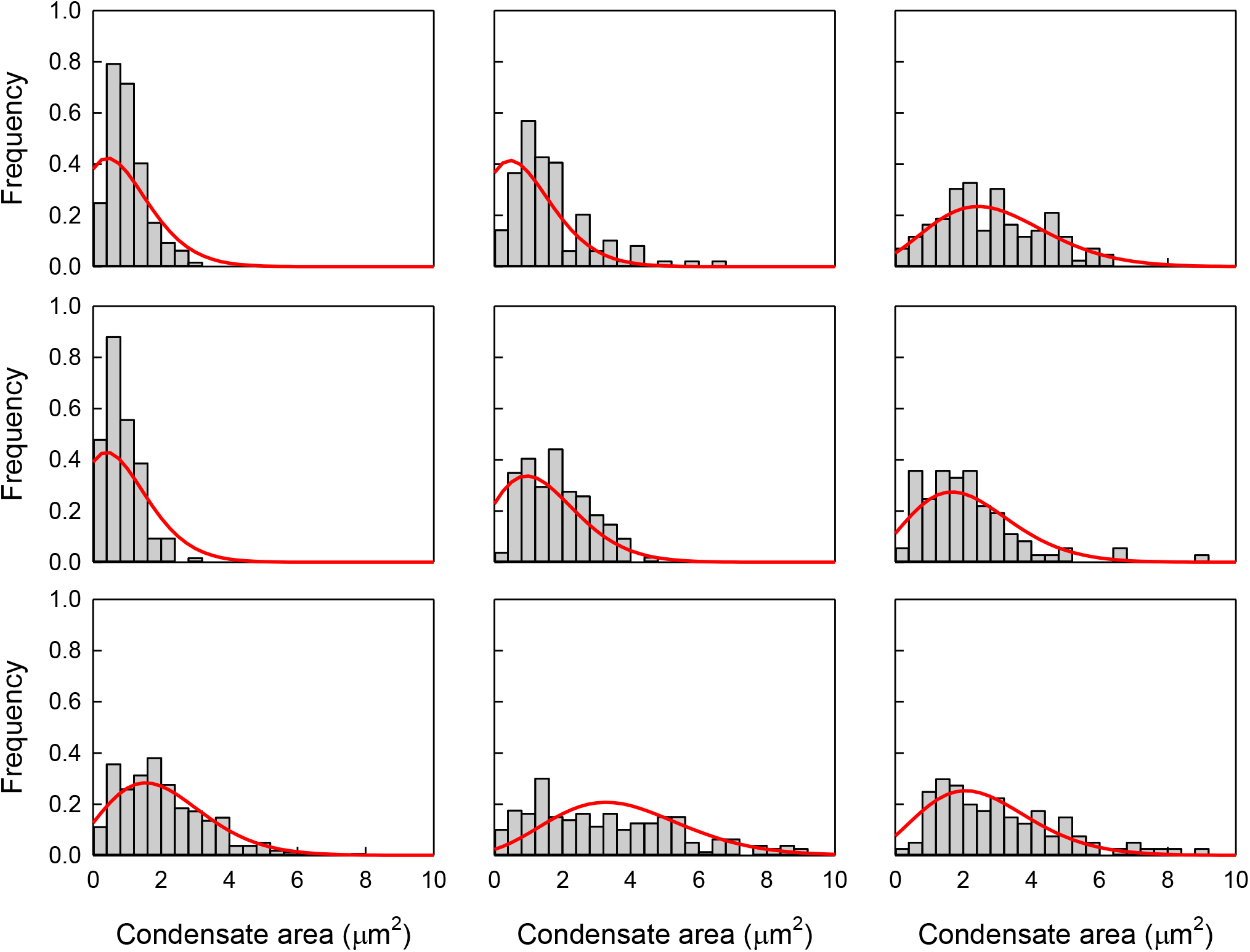
Condensate size distribution over time. Top row: polyA_10_ and [RGRGG]_4_ around (from left) 4:15, 11:05 and 20:56 minutes after mixing. Middle row: polyA_10_ and [RGRGG]_10_ around 3:23, 10:50 and 20:44 minutes after mixing. Bottom row: polyA_20_ and [RGRGG]_4_ around 3:32, 11:54 and 21:11 minutes after mixing. The bin width is 0.4 μm^2^. The red lines fit a Poisson distribution. Concentrations are the same as in Fig. 1.

While there was qualitatively good agreement between initial computer predictions and experiments, we considered how initial simulations using the COCOMO model may be improved to match experiments quantitatively. The model contains various parameters for different types of interactions, which have been calibrated using protein/RNA polymer properties data and condensation measurements from the literature. A straightforward modification would have been to increase the strength of the RNA-peptide cation-π interactions, as it makes phase separation more likely and would decrease the length threshold, similar to experiments. However, we also noted that the density in the condensed phase was extremely high^26-27^, suggesting there would be little to no water within them under experimental conditions^28-29^ (**Fig. S3**). Since the experimentally observed condensates showed spherical droplets exhibiting liquid-like growth over time, such high densities may be unexpected. Increasing the size of the beads, described via the σ_i_ parameter, by 20% decreased the density commensurately (**Fig. S3**). At the same time, increasing the size of the beads decreased the minimum lengths for condensation, in quantitative agreement with experiments. This modification to the model is referred as COCOMO 1.2σ and was employed in the rest of the simulations in this work.

**Fig. 1** shows that the phase boundary for condensation is anticorrelated in length of the two components; as polyA_N_ increases in length, the minimum peptide length required decreases while keeping the concentration of residues (adenine, arginine and glycine) the same. This suggests that the length-dependent condensation is primarily determined by entropy. This idea has been mentioned previously for other protein-RNA phase separating system^21^ and is supported by results from the CG model. **Fig. S4** shows that the radial distribution functions (RDFs) between residues in the condensates are not substantially changed by length. In other words, condensates of longer polymers are not more tightly bound than condensates of shorter ones.

Possible sources of entropy changes between disperse and condensed phases involve the loss of translational freedom, differences in conformational entropy and differences in mixing entropy (the peptide/RNA composition is different in the condensates than in the disperse phase). Based on the simulations, condensates have similar compositions of peptides vs. RNA as in the initial disperse systems (see **Table S5**). Therefore, the change in mixing entropy is small. To consider changes in conformational entropy, we examined the radii of gyration for each polymer combination in the condensed and disperse (before condensation is observed) phases. **Fig. S5** shows that they are indistinguishable for the measured lengths. Only for very long RNA and peptides are there more notable differences in radii of gyration, indicating that they become more extended within the condensate. More extended conformational ensembles in the condensed phase have been related to enhanced phase separation with disordered proteins, by reducing steric hindrance and therefore maximizing intermolecular contacts ^30^. Additionally, **Fig. S6** shows that the probability of intrachain distances between residue 1 and 5 in peptides and RNA is unchanged with polymer length and between the dilute and condensed phases. We therefore conclude that the conformational entropy of individual polymers is unchanged between phases. The only remaining source of entropy is therefore due to the confinement of individual polymers within the condensate.

To confirm this hypothesis, we built a thermodynamic model using details obtained from the CG simulation (see SI for details). The model focuses on estimating the stability of the condensates from the free energy based on enthalpy-entropy decomposition. The model holds the residue concentration constant and does not consider the conditions for phase coexistence required for condensation to be observed. To find the enthalpy, *h*, for each residue pair (i.e. adenine-adenine, adenine-arginine, adenine-glycine, arginine-arginine, arginine-glycine, and glycine-glycine) in the cluster, the RDF from the CG simulations is convolved with the CG potential, integrated and multiplied by the number density of residues. Relevant here is the change in enthalpy from the disperse phase to the condensed phase. The enthalpy in the disperse phase is approximately zero when only intrachain polymer interactions are considered. This assumes that there are no RNA-peptide interactions. In reality, some interactions occur in the disperse phase giving rise to a small, negative enthalpy contribution. However, we could only reliably estimate the interchain interaction contribution to the enthalpy in the disperse phase for some systems at the polymer length limits where condensate formation took long enough to collect reliable statistics from the initial disperse phase. In those cases, the total disperse phase enthalpy was 5-10% of the condensate enthalpy. In the other cases where condensation occurs quickly, it is not clear how to define the disperse phase and obtain RDFs. To circumvent these issues, we postulate a reference disperse state where there are no interchain polymer interactions. In that case, it is sufficient to calculate just the condensate enthalpy alone.

The change in entropy is estimated from the ratio of the accessible volume in the condensate to the volume of the box in the simulations (**Eq. S10**) The volume accessible to a polymer in the condensate was estimated from the molecular volume of the polymer with the argument that the free space inside the dense condensates is too fragmented for a given polymer to fit elsewhere and that the accessible volume is therefore just the volume already occupied by the polymer. It is important to note that while enthalpy contributions are estimated per residue, entropy is calculated per polymer since it is the translational freedom of each polymer that is being restricted. The additional restriction of rotational degrees of freedom was not considered in the analysis. Fig. 3 shows the resulting enthalpy, entropy and total free energy change between the dilute and condensed phase for each polymer mixture of polyA_5,10,20_ and [RGRGG]_1,2002C3,4,6,8,10_. With increasing length *ΔG* becomes negative at the simulated phase boundary.

**Figure 3.**
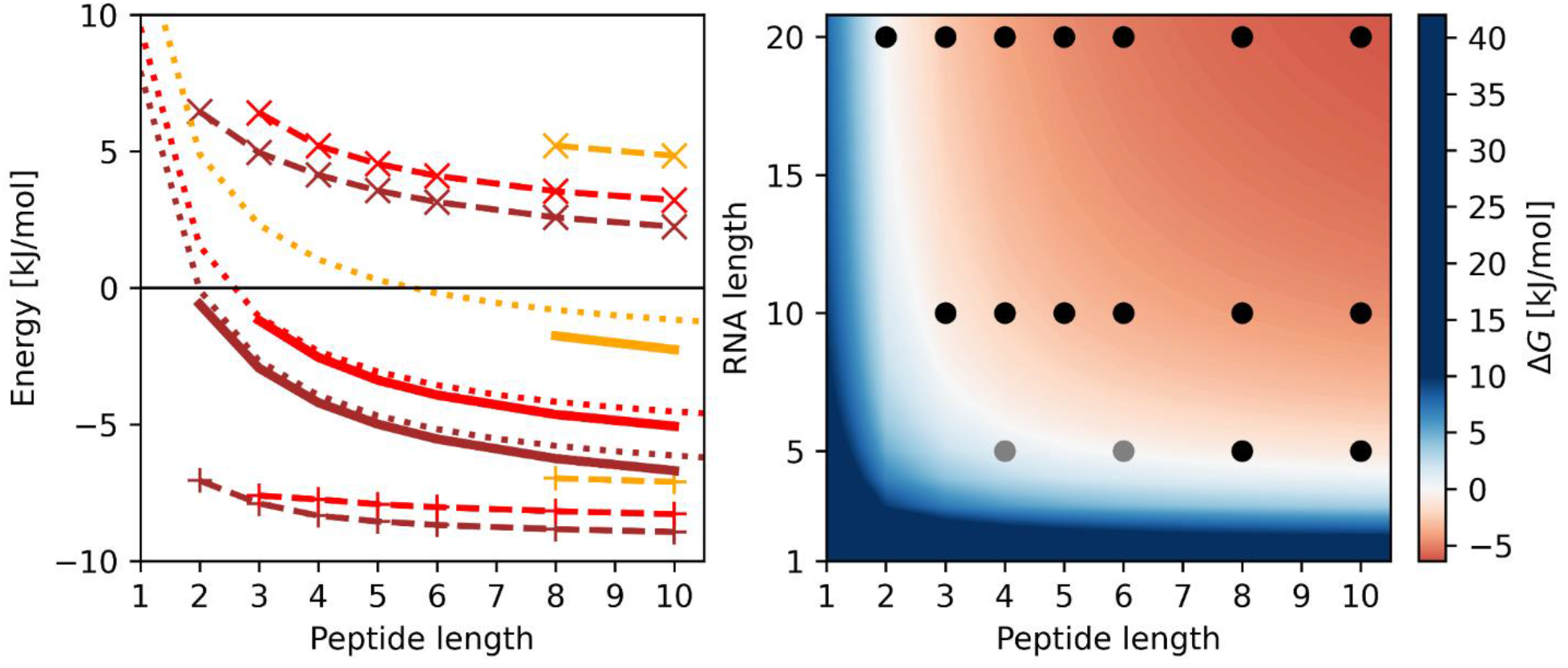
Energetic analysis of peptide-RNA condensates based on enthalpy-entropy composition. Condensate enthalpies according to **Eq. S3** (dashed lines with ‘+’), entropies (*-TΔS* at 300 K) according to **Eq. S9** (dashed lines with ‘x’), and total free energies according to **Eq. S2** (solid lines) are shown on the left as a function of peptide length with different RNA (polyA_20_: brown, polyA_10_: red, polyA_5_: orange). Energies were estimated by averaging over five replicate simulations. The statistical errors of the mean are less than 0.1 kJ/mol and are not shown. Dotted lines reflect total free energy estimates using densities and RDFs based on polyA_20_/[RGRGG]_3_ but renormalized to reflect clusters with a 50 nm radius in a box size with a length of 300 nm. The contour plot on the right shows the total free energies as a function of peptide and RNA length obtained with the same parameters. Dots indicate combinations of peptide/RNA for which condensates were observed experimentally (black and grey) and in the simulations (black only).

The experiments and simulations indicate minimum peptide and RNA lengths for condensation. However, we found that shorter peptides (i.e. [RGRGG]_1_) may participate in condensates once they are formed by longer peptides. This led us to speculate that shorter polymers may be able to compensate when the concentration of a longer peptide was too low to observe condensation. To demonstrate this phenomenon, we reduced the concentration of [RGRGG]_2_ in a mixture with polyA_20_ until condensation was lost, between 0.4 and 0.5 mg/mL. Then [RGRGG]_1_ was added until phase separation was recovered, between 0.6 and 0.7 mg/mL. This observation was confirmed by the CG model: the threshold for [RGRGG]_2_ condensation was ∼0.55 mg/mL and for [RGRGG]_1_ phase separation recovery was ∼1.8 mg/mL (see **Fig. 4** and **S7**). We also showed that [RGRGG]_1_ alone, even at high concentrations, cannot induce phase separation (**Fig. S8**).

**Figure 4.**
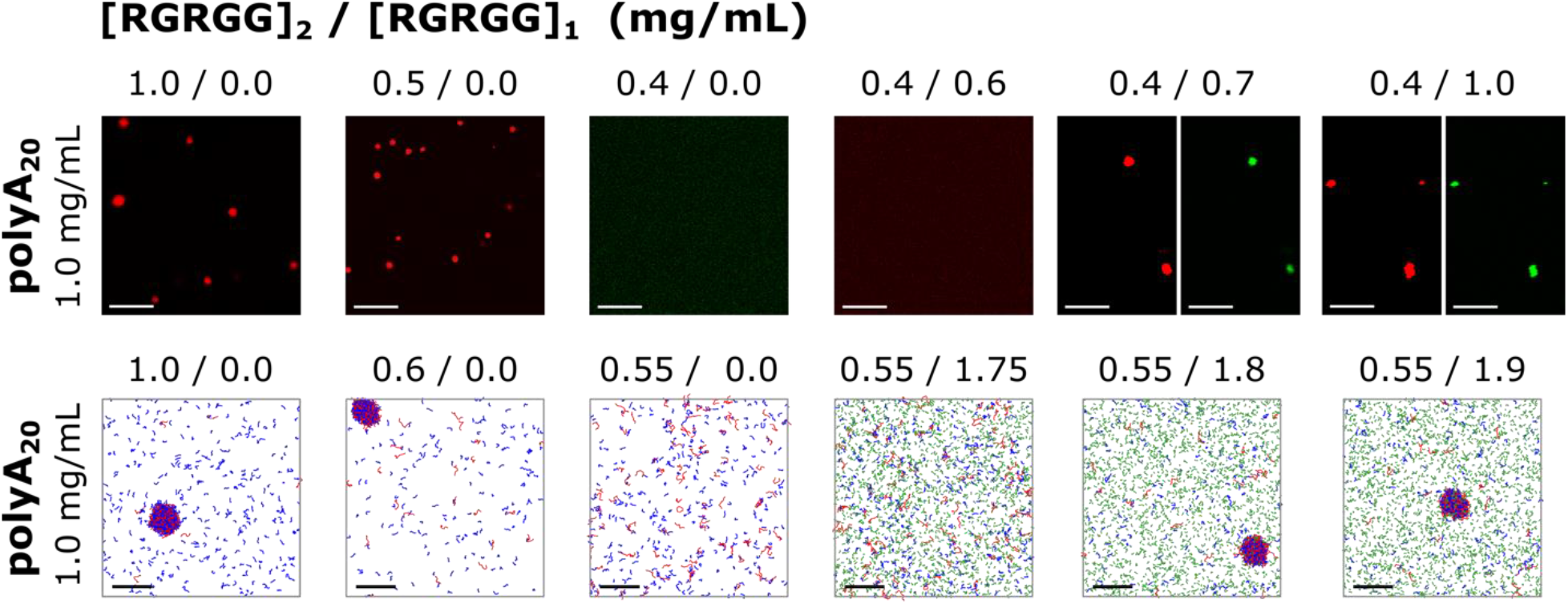
Phase separation recovery with small peptides. Experimental (upper panels) and simulation using COCOMO 1.2σ (lower panels) results show the loss of LLPS when lowering [RGRGG]_2_ concentration below a certain threshold and its recovery when adding enough [RGRGG]_1_. polyA_20_ was kept at 1 mg/mL for these assays. The red condensates in the upper panels show the fluorescence of [RGRGG]_2_-Cy3 and the green condensates show the fluorescence of [RGRGG]_1_-Cy5, indicating coexistence in the condensates. The final frames of the trajectories are shown in the lower panels for each simulated system with polyA_20_, [RGRGG]_1_, and [RGRGG]_2_ colored in red, green, and blue, respectively. Scale bars represent 10 μm in experimental panels and 20 nm in the simulation panels.

## CONCLUSION

We have shown systematically the effect of length on condensation of positively charged peptides and negatively charged RNA. We observed condensation for combinations of relatively short peptides and RNA, but experiments and simulations show there does exist a lower limit in terms of RNA or peptide lengths for the systems studied here. The essential driving force for condensation is electrostatic attraction between RNA and arginine residues, counteracted by the entropic cost of condensation. The key reason for the observed length dependence is that the enthalpy of condensation scales with the number of charged units whereas entropy scales with the number of polymers. We expect that with peptides with lower charge density, e.g., arginine residues spaced more widely, longer peptides and/or longer RNA would be required for phase separation to be observed. On the other hand, systems with higher charge density have been proven to enable phase separation even with single nucleotides, such as UDP (−3 charge) or UTP (−4 charge), and polyR_10_ as observed in other work^18^. Our study furthermore demonstrates that yet shorter polymers may participate in condensates as clients or as partial drivers of condensation together with longer polymers.

The remarkable agreement between experiment and CG simulations suggests that the computational approach could be extended to other sequences and systems as well as other dimensions in phase space, such as concentration or peptide content. The parameterization of the CG model^20^ explored a wide range of sequences and concentrations and accounts, for example, for the higher condensation propensity of arginine with nucleotides than lysine, as was observed by Fisher and Elbaum–Garfinkle^18^. However, the relative simplicity of the CG model neglects counterion effects that are known to be important factors during condensation^31-32^ and does not consider partial secondary structures that are present in many IDPs. Extending the model to explore the importance of these factors will be the subject of future studies.

LLPS has practical applications for inducing high concentration phases of certain biomolecules, the work here illustrates a quantitative framework for predicting the system components necessary for LLPS. On the other hand, this work demonstrates that a wide range of peptides and RNA can lead to LLPS. In the biological context, this means that many biomolecules may drive and/or participate in condensate formation in a dynamic manner as cellular concentrations of peptides and RNA fluctuate.

## METHODS

### Experimental

Liquid-liquid phase separation was studied for short proteins and RNA. The peptides [RGRGG]_1,2,3,4,6,8,10_ and Cy5-labeled [RGRGG]_1_ were obtained from Bio-Synthesis, Inc. RNA polyA_5,10,20_ and Cy3-labeled polyA_10,20_ were obtained from Horizon Discovery. These constructs were used without modification and dissolved in a 20 mM sodium phosphate buffer. Mixtures of unlabeled RNA and peptide were created at concentrations of 1 mg/ml, except where noted, along with 5μM fluorescently labeled samples.

A Nikon A1 Rsi Confocal Laser Scanning Microscope configured on an automated Nikon Eclipse Ti inverted microscope (Nikon Instruments, Inc.) equipped with a 100x Plan Apo TIRF oil objective (NA 1.45) was used to capture the confocal images at 100x objective magnification and PMT (photomultiplier tube) detector set to 31 HV. The Cy3 and Cy5 fluorescence were excited using a diode laser at 561 nm and 647 nm and recorded through 595/50 nm and 700/75 nm band pass emission filters respectively. Image acquisition was performed using the Nikon NIS Elements software (version 5.21.03). Transmitted light images were recorded using Differential Interference Contrast (DIC) optics at 561 nm.

Confocal images were generally obtained within 3 minutes of mixing for all combinations of RNA and peptide. To study the growth of condensates with time, 10 images were obtained during a time interval of ∼2 minutes for each of the following: right after mixing, and after 10 and 20 minutes of mixing. Size distribution analysis of condensates was performed using ImageJ software.

Brightfield microscopy was used to identify the effect of Cy3 on condensate formation. Images were obtained using an AmScope compound microscope equipped with a 10x objective (NA 0.25).

### Coarse-Grained Simulations

Initial observations of length-dependent cluster formation were done with simulations using the COCOMO model^20^ (Figure S1). To improve agreement with experimental data and reduce unexpectedly high polymer densities in the condensates, the short-range interaction parameter σ was increased 1.2 times for all residues and nucleotides. This modified version of the model is referred to as COCOMO 1.2σ. The model is described in more detail in the SI.

### Energetic analysis of peptide-RNA phase separation

Following an analytical treatment introduced by Dutagaci et al.^4^, the residue-level enthalpies and polymer-level entropies are calculated from the coarse-grained simulations. Details are given in the SI.

## ASSOCIATED CONTENT

### Supporting Information

Description of computational methods, tables and figures.

## Author Contributions

The manuscript was written through contributions of all authors. / All authors have given approval to the final version of the manuscript.

## ACKNOWLEDGMENT

Experimental data was collected at the Michigan State University Center for Advanced Microscopy.

## ABBREVIATIONS

CG: coarse-grained;
LLPS: liquid-liquid phase separation

## TOC graphic

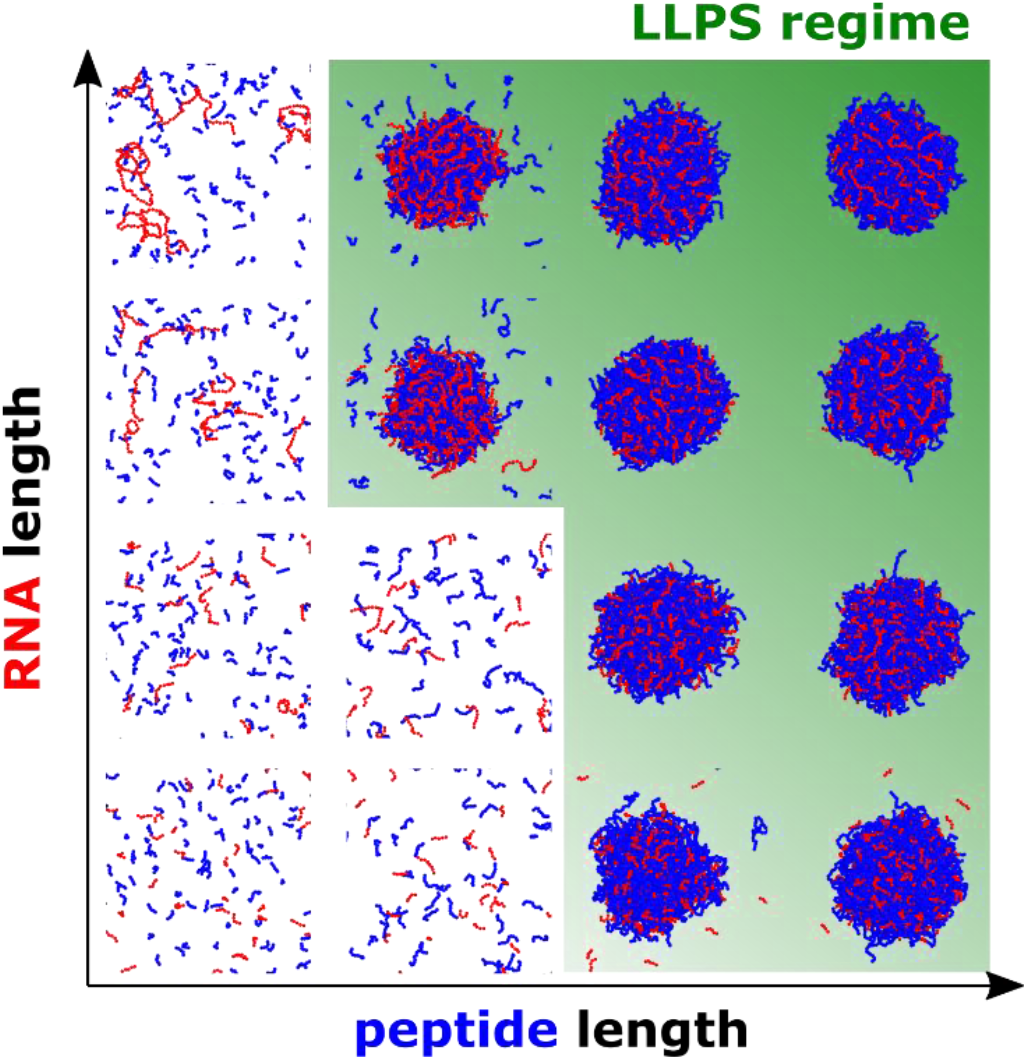

