## Supplemental Information for "The Effect of Polymer Length in Phase Separation"

*Coarse-grained simulations*

In our model each residue, protein or RNA, is represented as a spherical bead. The force field is given by:

$$U_{total} = \sum_{i=1}^{N-1} \frac{1}{2} k_{bond} \left(l_{i,i+1} - l_0\right)^2 + \sum_{i=1}^{N-2} \frac{1}{2} k_{angle} \left(\theta_{i,i+1,i+2} - \theta_0\right)^2$$

$$+ \sum_{i,j} 4(\varepsilon + \varepsilon_{cation-\pi}) \left( \left(\frac{\sigma_{i,j}}{r_{i,j}}\right)^{10} - \left(\frac{\sigma_{i,j}}{r_{i,j}}\right)^{5} \right) + \sum_{i,j} \frac{(A_{i,j} + A0_{i,j})}{r_{i,j}} e^{-\frac{r_{i,j}}{\kappa}} \quad \text{(S1)}$$

where bonded interactions are described using: $l$, distance between two neighboring residues; $k_{bond}$, spring constant equal to 4184 kJ/mol·nm$^2$; $l_0$, the equilibrium bond length equals 0.38 nm for proteins and 0.50 nm for RNA; $\theta$ the angle between three beads with an angle constant, $k_{angle}$, of 4.184 kJ/mol·rad$^2$ and equilibrium angle, $\theta_0$ the is 180°. Non-bonded interaction parameters were: $r_{i,j}$, inter-particle distance, $\sigma_{i,j}$, determined from the radius of a sphere of equivalent volume of a given residue, ε is set for polar and non-polar protein residues, and nucleotides. Cation-π interaction were added using a $\varepsilon_{cation-\pi}$ parameter set to 0.3 kJ/mol for R/K – F/Y/W and 0.2 kJ/mol for R/K – nucleotides pairwise interactions; $A_{ij}$ describes long-range interactions and for any residue $A_i = \text{sign}(q_i)\sqrt{0.75|q_i|}$[1] with $q_i$ being the residue charge, $A0_{i,j} = A0_i + A0_j$ describes the repulsion between polar residues due to solvation. Finally, $\kappa$ was set to 1 nm, what corresponds to an ionic strength of ~100 mM. **Table S1** reports all residue-specific parameters with the column specifying the σ values employed in the simulation with COCOMO and COCOMO 1.2σ.

Molecular dynamics simulations were performed using OpenMM 7.7.0[2]. Langevin dynamics was applied with a friction coefficient of 0.01 ps$^{-1}$, with temperature kept at 298 K. Systems were set up by randomly place enough copies of proteins and RNA in a 100 nm side box to fulfill a final concentration of 1 mg/mL of both molecules (**Tables S2-4**). Systems were initially minimized using 5,000 steps of steepest descent followed by 20,000 steps of MD with a time-step of 0.01 ps. Production runs were with a time step of 0.02 ps. Non-bonded interactions were truncated at a cutoff distance of 3 nm and calculated using periodic boundary conditions. Residues separated by one bond were excluded from non-bonded interactions. All systems were simulated for 20 μs with five replicates each. Coordinates are saved every 500 ps. The exact composition of the simulated systems using COCOMO and COCOMO 1.2σ are listed in Table S2 and Tables S3-4, respectively.

All figures from simulations were rendered in PyMOL (Schrodinger, LLC. The PyMOL Molecular Graphics System, Version 2.5.2 2021).

*Energetic analysis of peptide-RNA phase separation*

We followed an analytical treatment introduced earlier[1] to better understand the polymer length dependence in the peptide-RNA condensates described here via experiments and simulations.

To describe the polymer mixtures, we consider multi-component systems that are described either at the polymer level (for estimating entropy) or at the residue level (for estimating enthalpies). We use the term 'residue' to mean either an amino acid in a peptide or a nucleotide in RNA. At the molecule level, there are P (peptide) and N (RNA) particles present in total concentrations of $c_N$ and $c_P$. At the residue level, there are R (Arg) and G (Gly) amino acid residues and A (Ade) nucleotides at total concentrations of $c_R$, $c_G$, and $c_A$.

The total system volume is $V$. When phase separation occurs, a high-density condensate of volume $V_c$ is formed. The condensates we observed in experiments and simulations are spherical in shape and the size and volume of the condensate are therefore defined by the condensate radius $r_c$. In the simulations, the radius is determined as the distance from the center of the condensate where the density drops to half of the density at the center of the condensate.

The concentrations of the polymer particles in the condensed phase are denoted with an additional subscript c: $c_{P,c}$, $c_{N,c}$, $c_{R,c}$, $c_{R,c}$, $c_{G,c}$, and $c_{A,c}$. From the concentrations, number densities are obtained according to $\rho = \frac{c}{\text{mM}} \cdot \frac{N_A}{10^{27}\text{nm}^3}$.

Phase separation requires that the formation of the condensate is favorable according to free energy relative to the initial disperse phase. Phase coexistence of a given component furthermore requires that the chemical potentials in the condensed and dilute phases are equal to each other. In principle, both components (i.e. peptide and RNA), either one, or neither component may be in coexistence depending on the system composition and overall conditions. An accurate analytical treatment of phase coexistence for the peptide-RNA mixtures described here under consideration of their polymer properties and additional factors such as the role of ions is challenging and will be left for future work. Instead, we focus here on the simpler analysis of just the free energy of the condensate to determine under which conditions condensate formation itself is favorable.

The free energy inside a condensate is estimated from an enthalpy-entropy decomposition according to:

$$\Delta G = \Delta H - T\Delta S \quad \text{(S2)}$$

The enthalpy (per volume) is estimated based on residue-wise enthalpies $h$ and residue densities inside the condensate. As discussed in the main text, the enthalpy of the disperse reference state is neglected as it is essentially zero under the assumption that there are no interchain interactions.

$$\Delta H = \rho_{R,c} \cdot \Delta h_{R,c} + \rho_{G,c} \cdot \Delta h_{G,c} + \rho_{A,c} \cdot \Delta h_{A,c} \quad \text{(S3)}$$

Residue number densities were extracted from the simulations (Figure S3).

Each residue-wise enthalpy was calculated as a sum of pair-wise interactions with other residues in the condensates as well as self-interactions:

$$\Delta h_{R,c} = \frac{1}{2}(\Delta h_{RR,c} + \Delta h_{RG,c} + \Delta h_{RA,c}) \quad \text{(S4)}$$

$$\Delta h_{G,c} = \frac{1}{2}(\Delta h_{GR,c} + \Delta h_{GG,c} + \Delta h_{GA,c}) \quad \text{(S5)}$$

$$\Delta h_{A,c} = \frac{1}{2}(\Delta h_{AR,c} + \Delta h_{AG,c} + \Delta h_{AA,c}) \quad \text{(S6)}$$

Where the factor 1/2 corrects for double-counted interactions.

Each pair-wise contribution was then calculated from the convolution of pair-wise radial distribution functions with the interaction potential according to the coarse-grained interaction potential (Eq. S7) that depends on the parameters for the interacting residues as described above. Accordingly, the enthalpy contribution for any two residue types X and Y is calculated as:

$$\Delta h_{XY,c} = \rho_{X,c} \int_V \hat{g}_{XY,d}(r) U_{XY}(r) d^3r = 4\pi \rho_{X,c} \int_0^{r_{max}} \hat{g}_{XY,d}(r) U_{XY}(r) r^2 dr \quad \text{(S7)}$$

The radial distribution functions were extracted from the CG simulations (Figure S4). Because of the finite-size of the condensates, the extracted *g(r)* functions decrease with increasing radius according to [3]:

$$g(r) = \left[1 - \frac{3}{2}\left(\frac{r}{d}\right) + \frac{1}{2}\left(\frac{r}{d}\right)^3\right] g_\infty(r) \quad \text{(S8)}$$

when determined from particles distributed homogeneously in a finite size sphere of radius *d*.

To compare finite-size condensates with theoretically infinitely large condensates, we considered both the original *g(r)* functions extracted from the simulations as well as normalized functions according to Eq. S8 to obtain $g(r_{cut})$=1 at a given cutoff distance of $r_{cut}$ (set to 12 nm) and a constant value of 1 for all larger radii. Such normalized g(r) functions could then be rescaled again via Eq. S8 to reflect theoretical distribution functions for larger condensates such as those observed experimentally.

The upper integration limit $r_{max}$ was set to 15 nm for all interactions. At that radius and above, the interaction potential *U(r)* becomes negligible.

Bonded interactions were not considered in this analysis, but self-interactions between residues were omitted for next-neighbor bonded residues within the same polymer. Otherwise, intra- and inter-chain interactions were considered equivalently in the estimation of the condensate enthalpy.

The entropy term was calculated based on molar entropies corresponding to the loss of translational freedom in the condensate:

$$\Delta S = \rho_{P,c} \cdot \Delta s_{P,c} + \rho_{N,c} \cdot \Delta s_{N,c} \tag{S9}$$

An important point to reiterate here is that while enthalpy depends on residue interactions, and therefore scales with residue densities, the loss of translational entropy applies to each molecule no matter how large and therefore scales with molecule number densities.

The molar entropies are then calculated form the ratio of available volume in the condensed phase to the system volume:

$$\Delta s_{X,c} = R\log\left(\frac{V_{X,c}}{V}\right) \tag{S10}$$

where R is the universal gas constant and X is either P or N for peptides or RNA, respectively.

The system volume may be given as the simulation box size (*i.e.* $(100\text{ nm})^3$), or larger when considering the macroscopic systems studied in experiments. To estimate the accessible volume inside the condensate, we assumed that it is equal to the molecular volume of a given polymer, *i.e.* we assume that the high density in the condensate effectively provides such confinement and polymer entanglement that most of the translational motion is 'in place' around the position of a polymer inside the condensate. Both simulations and experiments showing liquid-like behavior for the condensates suggest that this is not entirely true, but it is likely a better approximation than assuming that all free volume inside the condensate that is not occupied by a polymer is accessible.

This analysis neglects other contributions to entropy such as mixing entropy as the composition inside the condensate and outside were similar based on the simulations. We also assume that conformational entropy per polymer segment is not significantly different as a function of polymer length and between polymers in the condensate and in the disperse phase as suggested by the simulations (Figure S5-6).

A program implementing this model is available at http://github.com/feiglab/peprnallps.

**Table S1. Residue-specific parameters used in COCOMO.**

| Residue | Mass (amu) | COCOMO $\sigma_i$ (nm)† | COCOMO 1.2σ $\sigma_i$ (nm)† | charge | $A_i$‡ | $A0_i$ | ε (kJ/mol) |
|---|---|---|---|---|---|---|---|
| Ala | 71.08 | 0.253 | 0.304 | 0 | 0 | 0 | 0.40 |
| Arg | 157.20 | 0.318 | 0.381 | 1 | 0.87 | 0.05 | 0.41 |
| Asn | 114.10 | 0.281 | 0.337 | 0 | 0 | 0.05 | 0.41 |
| Asp | 114.08 | 0.277 | 0.333 | -1 | -0.87 | 0.05 | 0.41 |
| Cys | 103.14 | 0.269 | 0.323 | 0 | 0 | 0.05 | 0.41 |
| Gln | 128.13 | 0.295 | 0.354 | 0 | 0 | 0.05 | 0.41 |
| Glu | 128.11 | 0.292 | 0.351 | -1 | -0.87 | 0.05 | 0.41 |
| Gly | 57.05 | 0.233 | 0.280 | 0 | 0 | 0 | 0.40 |
| His | 137.14 | 0.297 | 0.357 | 0 | 0 | 0.05 | 0.41 |
| Ile | 113.16 | 0.299 | 0.359 | 0 | 0 | 0 | 0.40 |
| Leu | 113.16 | 0.300 | 0.360 | 0 | 0 | 0 | 0.40 |
| Lys | 129.18 | 0.306 | 0.368 | 1 | 0.87 | 0.05 | 0.41 |
| Met | 131.19 | 0.301 | 0.361 | 0 | 0 | 0 | 0.40 |
| Phe | 147.18 | 0.317 | 0.380 | 0 | 0 | 0 | 0.40 |
| Pro | 98.13 | 0.284 | 0.341 | 0 | 0 | 0 | 0.40 |
| Ser | 87.08 | 0.261 | 0.313 | 0 | 0 | 0.05 | 0.41 |
| Thr | 101.11 | 0.277 | 0.332 | 0 | 0 | 0.05 | 0.41 |
| Trp | 186.21 | 0.334 | 0.401 | 0 | 0 | 0 | 0.40 |
| Tyr | 71.08 | 0.322 | 0.386 | 0 | 0 | 0 | 0.40 |
| Val | 157.20 | 0.286 | 0.343 | 1 | 0 | 0 | 0.40 |
| Ade | 315.70 | 0.376 | 0.451 | -1 | -0.87 | 0.05 | 0.41 |
| Cyt | 305.20 | 0.366 | 0.439 | -1 | -0.87 | 0.05 | 0.41 |
| Gua | 345.20 | 0.379 | 0.455 | -1 | -0.87 | 0.05 | 0.41 |
| Ura | 305.16 | 0.364 | 0.437 | -1 | -0.87 | 0.05 | 0.41 |

† $\sigma_i = 2^{-1/6}\, r_i$, where $r_i$ is the radius of a sphere with equivalent volume of residue *i*.

‡ $A_i = \text{sign}(q_i)\sqrt{0.75|q_i|}$, where $q_i$ is the residue net charge of the residue *i*.

**Table S2.** Composition of protein – RNA systems for the length dependence simulations using COCOMO.

| System | conc (mM) | N chains | System | conc (mM) | N chains |
|---|---|---|---|---|---|
| polyA$_5$ / [RGRGG]$_1$ | 0.63 / 2.06 | 376 / 1240 | polyA$_{50}$ / [RGRGG]$_1$ | 0.06 / 2.06 | 38 / 1240 |
| polyA$_5$ / [RGRGG]$_2$ | 0.63 / 1.03 | 376 / 620 | polyA$_{50}$ / [RGRGG]$_2$ | 0.06 / 1.03 | 38 / 620 |
| polyA$_5$ / [RGRGG]$_3$ | 0.63 / 0.68 | 376 / 414 | polyA$_{50}$ / [RGRGG]$_3$ | 0.06 / 0.68 | 38 / 414 |
| polyA$_5$ / [RGRGG]$_4$ | 0.63 / 0.52 | 376 / 310 | polyA$_{50}$ / [RGRGG]$_4$ | 0.06 / 0.52 | 38 / 310 |
| polyA$_5$ / [RGRGG]$_5$ | 0.63 / 0.41 | 376 / 248 | polyA$_{50}$ / [RGRGG]$_5$ | 0.06 / 0.41 | 38 / 248 |
| polyA$_5$ / [RGRGG]$_6$ | 0.63 / 0.34 | 376 / 207 | polyA$_{50}$ / [RGRGG]$_6$ | 0.06 / 0.34 | 38 / 207 |
| polyA$_5$ / [RGRGG]$_8$ | 0.63 / 0.26 | 376 / 155 | polyA$_{50}$ / [RGRGG]$_8$ | 0.06 / 0.26 | 38 / 155 |
| polyA$_5$ / [RGRGG]$_{10}$ | 0.63 / 0.21 | 376 / 124 | polyA$_{50}$ / [RGRGG]$_{10}$ | 0.06 / 0.21 | 38 / 124 |
| polyA$_5$ / [RGRGG]$_{15}$ | 0.63 / 0.14 | 376 / 83 | polyA$_{50}$ / [RGRGG]$_{15}$ | 0.06 / 0.14 | 38 / 83 |
| polyA$_{10}$ / [RGRGG]$_1$ | 0.31 / 2.06 | 188 / 1240 | polyA$_{100}$ / [RGRGG]$_1$ | 0.03 / 2.06 | 19 / 1240 |
| polyA$_{10}$ / [RGRGG]$_2$ | 0.31 / 1.03 | 188 / 620 | polyA$_{100}$ / [RGRGG]$_2$ | 0.03 / 1.03 | 19 / 620 |
| polyA$_{10}$ / [RGRGG]$_3$ | 0.31 / 0.68 | 188 / 414 | polyA$_{100}$ / [RGRGG]$_3$ | 0.03 / 0.68 | 19 / 414 |
| polyA$_{10}$ / [RGRGG]$_4$ | 0.31 / 0.52 | 188 / 310 | polyA$_{100}$ / [RGRGG]$_4$ | 0.03 / 0.52 | 19 / 310 |
| polyA$_{10}$ / [RGRGG]$_5$ | 0.31 / 0.41 | 188 / 248 | polyA$_{100}$ / [RGRGG]$_5$ | 0.03 / 0.41 | 19 / 248 |
| polyA$_{10}$ / [RGRGG]$_6$ | 0.31 / 0.34 | 188 / 207 | polyA$_{100}$ / [RGRGG]$_6$ | 0.03 / 0.34 | 19 / 207 |
| polyA$_{10}$ / [RGRGG]$_8$ | 0.31 / 0.26 | 188 / 155 | polyA$_{100}$ / [RGRGG]$_8$ | 0.03 / 0.26 | 19 / 155 |
| polyA$_{10}$ / [RGRGG]$_{10}$ | 0.31 / 0.21 | 188 / 124 | polyA$_{100}$ / [RGRGG]$_{10}$ | 0.03 / 0.21 | 19 / 124 |
| polyA$_{10}$ / [RGRGG]$_{15}$ | 0.31 / 0.14 | 188 / 83 | polyA$_{100}$ / [RGRGG]$_{15}$ | 0.03 / 0.14 | 19 / 83 |
| polyA$_{20}$ / [RGRGG]$_1$ | 0.16 / 2.06 | 94 / 1240 | polyA$_{300}$ / [RGRGG]$_1$ | 0.01 / 2.06 | 6 / 1240 |
| polyA$_{20}$ / [RGRGG]$_2$ | 0.16 / 1.03 | 94 / 620 | polyA$_{300}$ / [RGRGG]$_2$ | 0.01 / 1.03 | 6 / 620 |
| polyA$_{20}$ / [RGRGG]$_3$ | 0.16 / 0.68 | 94 / 414 | polyA$_{300}$ / [RGRGG]$_3$ | 0.01 / 0.68 | 6 / 414 |
| polyA$_{20}$ / [RGRGG]$_4$ | 0.16 / 0.52 | 94 / 310 | polyA$_{300}$ / [RGRGG]$_4$ | 0.01 / 0.52 | 6 / 310 |
| polyA$_{20}$ / [RGRGG]$_5$ | 0.16 / 0.41 | 94 / 248 | polyA$_{300}$ / [RGRGG]$_5$ | 0.01 / 0.41 | 6 / 248 |
| polyA$_{20}$ / [RGRGG]$_6$ | 0.16 / 0.34 | 94 / 207 | polyA$_{300}$ / [RGRGG]$_6$ | 0.01 / 0.34 | 6 / 207 |
| polyA$_{20}$ / [RGRGG]$_8$ | 0.16 / 0.26 | 94 / 155 | polyA$_{300}$ / [RGRGG]$_8$ | 0.01 / 0.26 | 6 / 155 |
| polyA$_{20}$ / [RGRGG]$_{10}$ | 0.16 / 0.21 | 94 / 124 | polyA$_{300}$ / [RGRGG]$_{10}$ | 0.01 / 0.21 | 6 / 124 |
| polyA$_{20}$ / [RGRGG]$_{15}$ | 0.16 / 0.14 | 94 / 83 | polyA$_{300}$ / [RGRGG]$_{15}$ | 0.01 / 0.14 | 6 / 83 |
| polyA$_{30}$ / [RGRGG]$_1$ | 0.10 / 2.06 | 63 / 1240 | | | |
| polyA$_{30}$ / [RGRGG]$_2$ | 0.10 / 1.03 | 63 / 620 | | | |
| polyA$_{30}$ / [RGRGG]$_3$ | 0.10 / 0.68 | 63 / 414 | | | |
| polyA$_{30}$ / [RGRGG]$_4$ | 0.10 / 0.52 | 63 / 310 | | | |
| polyA$_{30}$ / [RGRGG]$_5$ | 0.10 / 0.41 | 63 / 248 | | | |
| polyA$_{30}$ / [RGRGG]$_6$ | 0.10 / 0.34 | 63 / 207 | | | |
| polyA$_{30}$ / [RGRGG]$_8$ | 0.10 / 0.26 | 63 / 155 | | | |
| polyA$_{30}$ / [RGRGG]$_{10}$ | 0.10 / 0.21 | 63 / 124 | | | |
| polyA$_{30}$ / [RGRGG]$_{15}$ | 0.10 / 0.14 | 63 / 83 | | | |

Initial concentrations of 1 mg/mL for both protein and RNA. All boxes were 100 nm side and simulation temperature 298 K.

**Table S3.** Composition of protein – RNA systems for the length-dependence simulations using COCOMO 1.2σ.

| System | conc (mM) | N chains | System | conc (mM) | N chains |
|---|---|---|---|---|---|
| polyA$_5$ / [RGRGG]$_1$ | 0.63 / 2.06 | 376 / 1240 | polyA$_{100}$ / [RGRGG]$_1$ | 0.03 / 2.06 | 19 / 1240 |
| polyA$_5$ / [RGRGG]$_2$ | 0.63 / 1.03 | 376 / 620 | polyA$_{100}$ / [RGRGG]$_2$ | 0.03 / 1.03 | 19 / 620 |
| polyA$_5$ / [RGRGG]$_3$ | 0.63 / 0.68 | 376 / 414 | polyA$_{100}$ / [RGRGG]$_3$ | 0.03 / 0.68 | 19 / 414 |
| polyA$_5$ / [RGRGG]$_4$ | 0.63 / 0.52 | 376 / 310 | polyA$_{100}$ / [RGRGG]$_4$ | 0.03 / 0.52 | 19 / 310 |
| polyA$_5$ / [RGRGG]$_5$ | 0.63 / 0.41 | 376 / 248 | polyA$_{100}$ / [RGRGG]$_5$ | 0.03 / 0.41 | 19 / 248 |
| polyA$_5$ / [RGRGG]$_6$ | 0.63 / 0.34 | 376 / 207 | polyA$_{100}$ / [RGRGG]$_6$ | 0.03 / 0.34 | 19 / 207 |
| polyA$_5$ / [RGRGG]$_8$ | 0.63 / 0.26 | 376 / 155 | polyA$_{100}$ / [RGRGG]$_8$ | 0.03 / 0.26 | 19 / 155 |
| polyA$_5$ / [RGRGG]$_{10}$ | 0.31 / 0.21 | 376 / 124 | polyA$_{100}$ / [RGRGG]$_{10}$ | 0.03 / 0.21 | 19 / 124 |
| polyA$_{10}$ / [RGRGG]$_1$ | 0.31 / 2.06 | 188 / 1240 | polyA$_{300}$ / [RGRGG]$_1$ | 0.01 / 2.06 | 6 / 1240 |
| polyA$_{10}$ / [RGRGG]$_2$ | 0.31 / 1.03 | 188 / 620 | polyA$_{300}$ / [RGRGG]$_2$ | 0.01 / 1.03 | 6 / 620 |
| polyA$_{10}$ / [RGRGG]$_3$ | 0.31 / 0.68 | 188 / 414 | polyA$_{300}$ / [RGRGG]$_3$ | 0.01 / 0.68 | 6 / 414 |
| polyA$_{10}$ / [RGRGG]$_4$ | 0.31 / 0.52 | 188 / 310 | polyA$_{300}$ / [RGRGG]$_4$ | 0.01 / 0.52 | 6 / 310 |
| polyA$_{10}$ / [RGRGG]$_5$ | 0.31 / 0.41 | 188 / 248 | polyA$_{300}$ / [RGRGG]$_5$ | 0.01 / 0.41 | 6 / 248 |
| polyA$_{10}$ / [RGRGG]$_6$ | 0.31 / 0.34 | 188 / 207 | polyA$_{300}$ / [RGRGG]$_6$ | 0.01 / 0.34 | 6 / 207 |
| polyA$_{10}$ / [RGRGG]$_8$ | 0.31 / 0.26 | 188 / 155 | polyA$_{300}$ / [RGRGG]$_8$ | 0.01 / 0.26 | 6 / 155 |
| polyA$_{10}$ / [RGRGG]$_{10}$ | 0.31 / 0.21 | 188 / 124 | polyA$_{300}$ / [RGRGG]$_{10}$ | 0.01 / 0.21 | 6 / 124 |
| polyA$_{20}$ / [RGRGG]$_1$ | 0.16 / 2.06 | 94 / 1240 | | | |
| polyA$_{20}$ / [RGRGG]$_2$ | 0.16 / 1.03 | 94 / 620 | | | |
| polyA$_{20}$ / [RGRGG]$_3$ | 0.16 / 0.68 | 94 / 414 | | | |
| polyA$_{20}$ / [RGRGG]$_4$ | 0.16 / 0.52 | 94 / 310 | | | |
| polyA$_{20}$ / [RGRGG]$_5$ | 0.16 / 0.41 | 94 / 248 | | | |
| polyA$_{20}$ / [RGRGG]$_6$ | 0.16 / 0.34 | 94 / 207 | | | |
| polyA$_{20}$ / [RGRGG]$_8$ | 0.16 / 0.26 | 94 / 155 | | | |
| polyA$_{20}$ / [RGRGG]$_{10}$ | 0.16 / 0.21 | 94 / 124 | | | |

Initial concentrations of 1 mg/mL for both protein and RNA. All boxes were 100 nm side and simulation temperature 298 K.

**Table S4.** Composition of protein – RNA systems for phase separation recovery with small peptides using COCOMO 1.2σ.

| System | conc (mg/mL) | conc (mM) | N chains |
|---|---|---|---|
| polyA$_{20}$ / [RGRGG]$_2$ / [RGRGG]$_1$ | 1.0 / 1.0 / 0 | 0.16 / 1.03 / 0 | 94 / 620 / 0 |
| polyA$_{20}$ / [RGRGG]$_2$ / [RGRGG]$_1$ | 1.0 / 0.6 / 0 | 0.16 / 0.62 / 0 | 94 / 372 / 0 |
| polyA$_{20}$ / [RGRGG]$_2$ / [RGRGG]$_1$ | 1.0 / 0.55 / 0 | 0.16 / 0.57 / 0 | 94 / 341 / 0 |
| polyA$_{20}$ / [RGRGG]$_2$ / [RGRGG]$_1$ | 1.0 / 0.55 / 1.75 | 0.16 / 0.57 / 3.6 | 94 / 341 / 2170 |
| polyA$_{20}$ / [RGRGG]$_2$ / [RGRGG]$_1$ | 1.0 / 0.55 / 1.8 | 0.16 / 0.57 / 3.7 | 94 / 341 / 2232 |
| polyA$_{20}$ / [RGRGG]$_2$ / [RGRGG]$_1$ | 1.0 / 0.55 / 1.9 | 0.16 / 0.57 / 3.9 | 94 / 341 / 2356 |
| polyA$_{20}$ / [RGRGG]$_2$ / [RGRGG]$_1$ | 1.0 / 0 / 1.8 | 0.16 / 0 / 3.7 | 94 / 0 / 2232 |

All boxes were 100 nm side and simulation temperature 298 K.

**Table S5.** Mixing ratios of number of RNA and peptide polymers in the system and within the condensates using COCOMO 1.2σ.

| System | RNA/peptide system | RNA/peptide condensate |
|---|---|---|
| polyA$_5$ / [RGRGG]$_8$ | 2.425806 | 1.880472 |
| polyA$_5$ / [RGRGG]$_{10}$ | 3.032258 | 2.308512 |
| polyA$_{10}$ / [RGRGG]$_3$ | 0.454106 | 0.649806 |
| polyA$_{10}$ / [RGRGG]$_4$ | 0.606452 | 0.734356 |
| polyA$_{10}$ / [RGRGG]$_5$ | 0.758065 | 0.829881 |
| polyA$_{10}$ / [RGRGG]$_6$ | 0.908213 | 0.940779 |
| polyA$_{10}$ / [RGRGG]$_8$ | 1.212903 | 1.214786 |
| polyA$_{10}$ / [RGRGG]$_{10}$ | 1.516129 | 1.513638 |
| polyA$_{20}$ / [RGRGG]$_2$ | 0.151613 | 0.312088 |
| polyA$_{20}$ / [RGRGG]$_3$ | 0.227053 | 0.332363 |
| polyA$_{20}$ / [RGRGG]$_4$ | 0.303226 | 0.371112 |
| polyA$_{20}$ / [RGRGG]$_5$ | 0.379032 | 0.418182 |
| polyA$_{20}$ / [RGRGG]$_6$ | 0.454106 | 0.473101 |
| polyA$_{20}$ / [RGRGG]$_8$ | 0.606452 | 0.608775 |
| polyA$_{20}$ / [RGRGG]$_{10}$ | 0.758065 | 0.758223 |

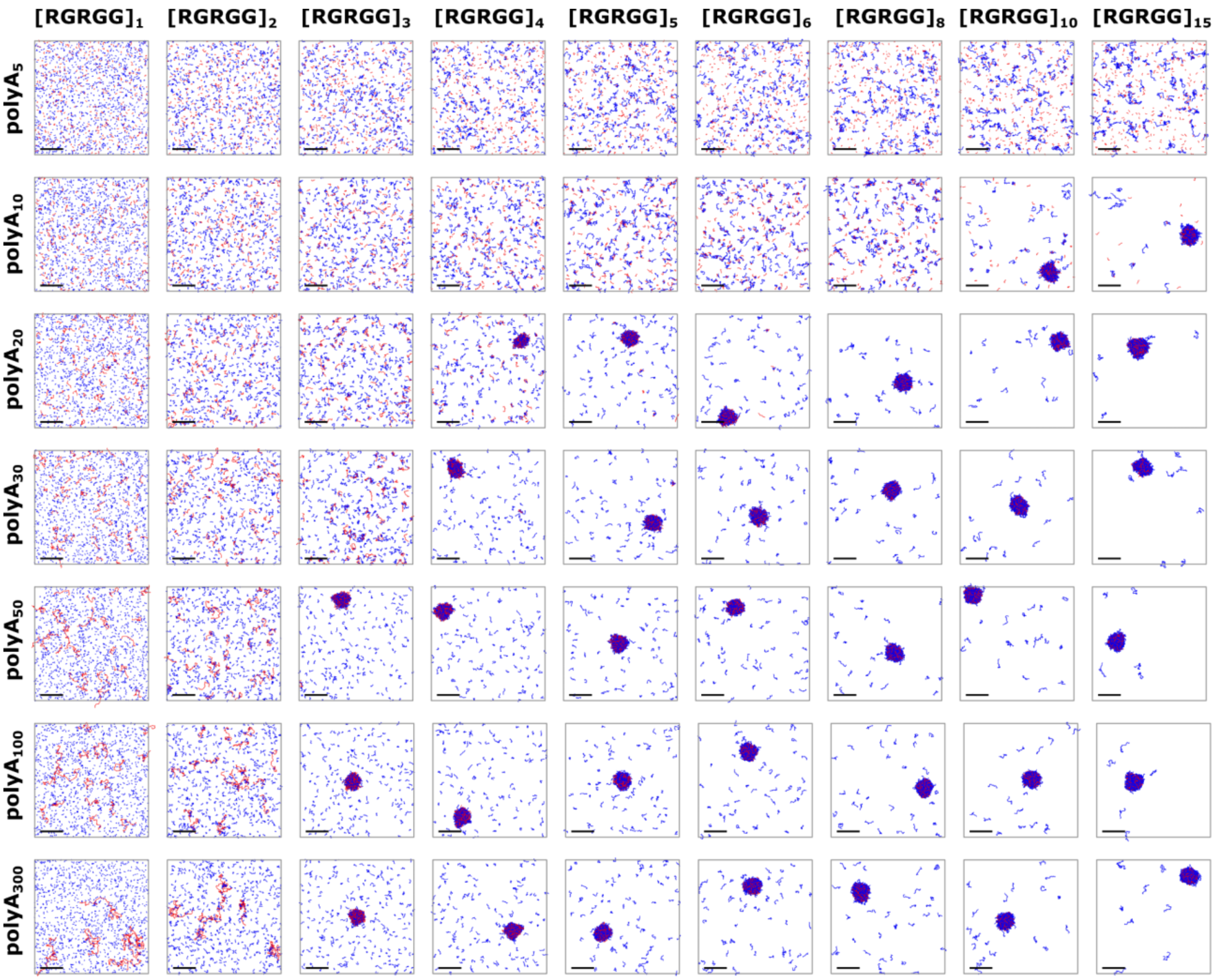


**Figure S1.** Length-dependent condensate formation of different protein/RNA systems. Simulations were performed using COCOMO without any modification. Final frame for each system's trajectory is shown with RNA and peptides colored in red and blue, respectively. Scale bars represent 20 nm.

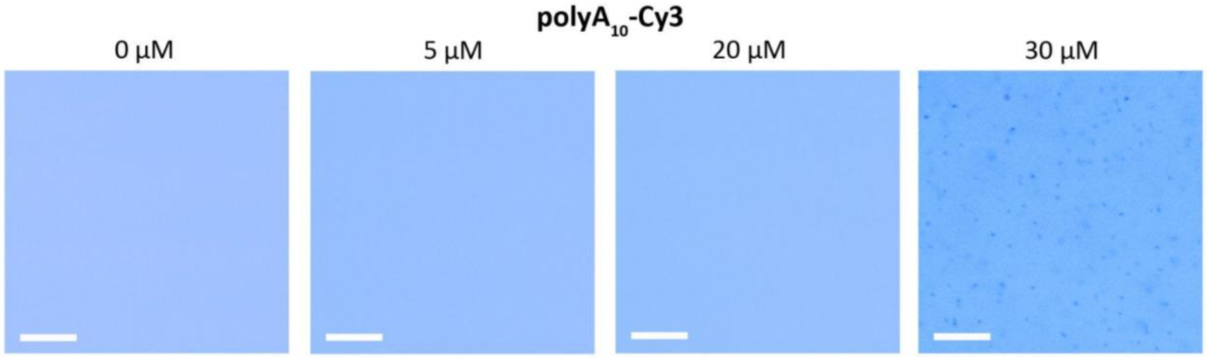


**Figure S2.** Inducement of condensates by Cy3. All samples contain polyA$_{10}$ and [RGRGG]$_2$ at the same total concentrations of 1mg/mL with polyA$_{10}$-Cy3 at 30 µM showing condensates and no condensates were observed at and below 20 µM. Scale bars represent 30 µm.

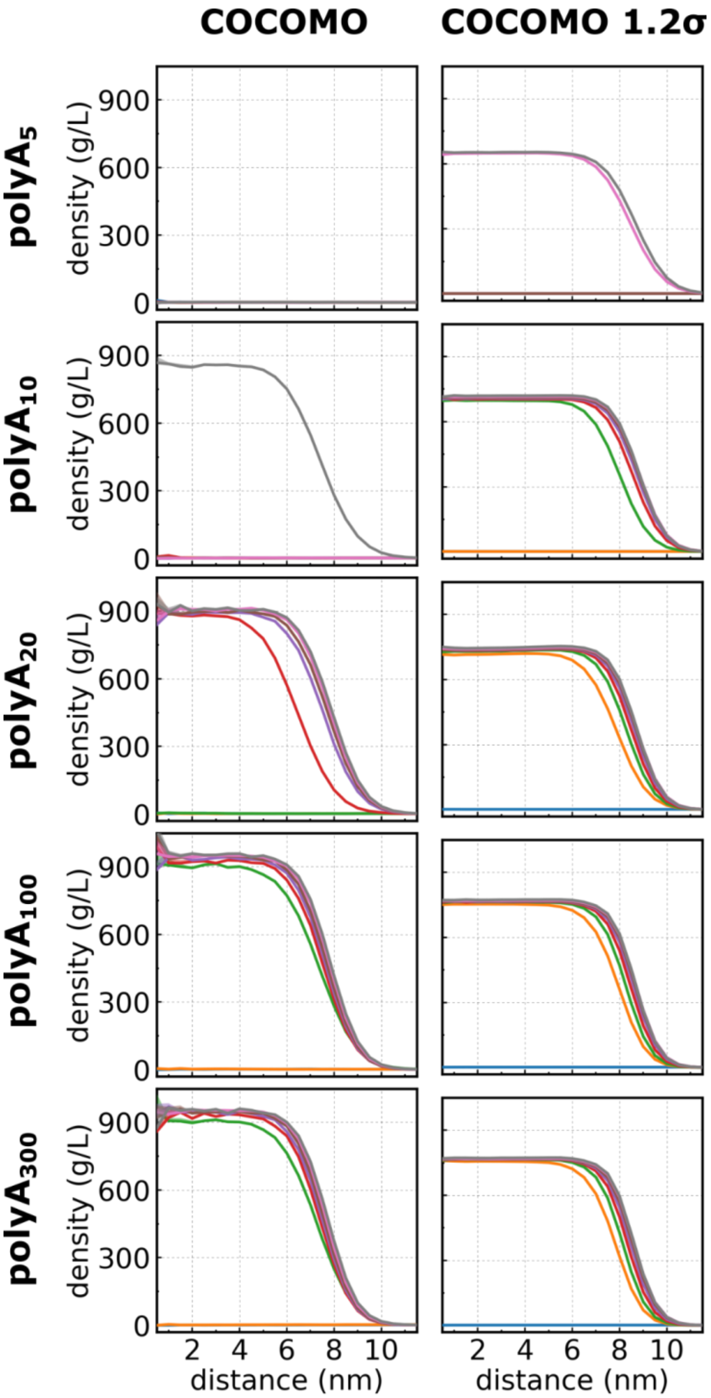


**Figure S3.** Cluster density in the simulation of different protein/RNA systems. Density ($\rho$) is shown as a function of distance, from the center of the condensate outwards. Results are shown column-wise for COCOMO and COCOMO - 1.2$\sigma$, and row-wise for different polyA$_N$ (N = 5, 10, 20, 100, and 300). Each trace shows a different [RGRGG]$_M$ peptide where M = 1, 2, 3, 4, 5, 6, 8, and 10 are in blue, orange, green, red, purple, brown, and pink, respectively. Flat density lines mean no clusters were formed in that particular system. Initial concentrations for these simulations were kept at 1 mg/mL for both protein and RNA.

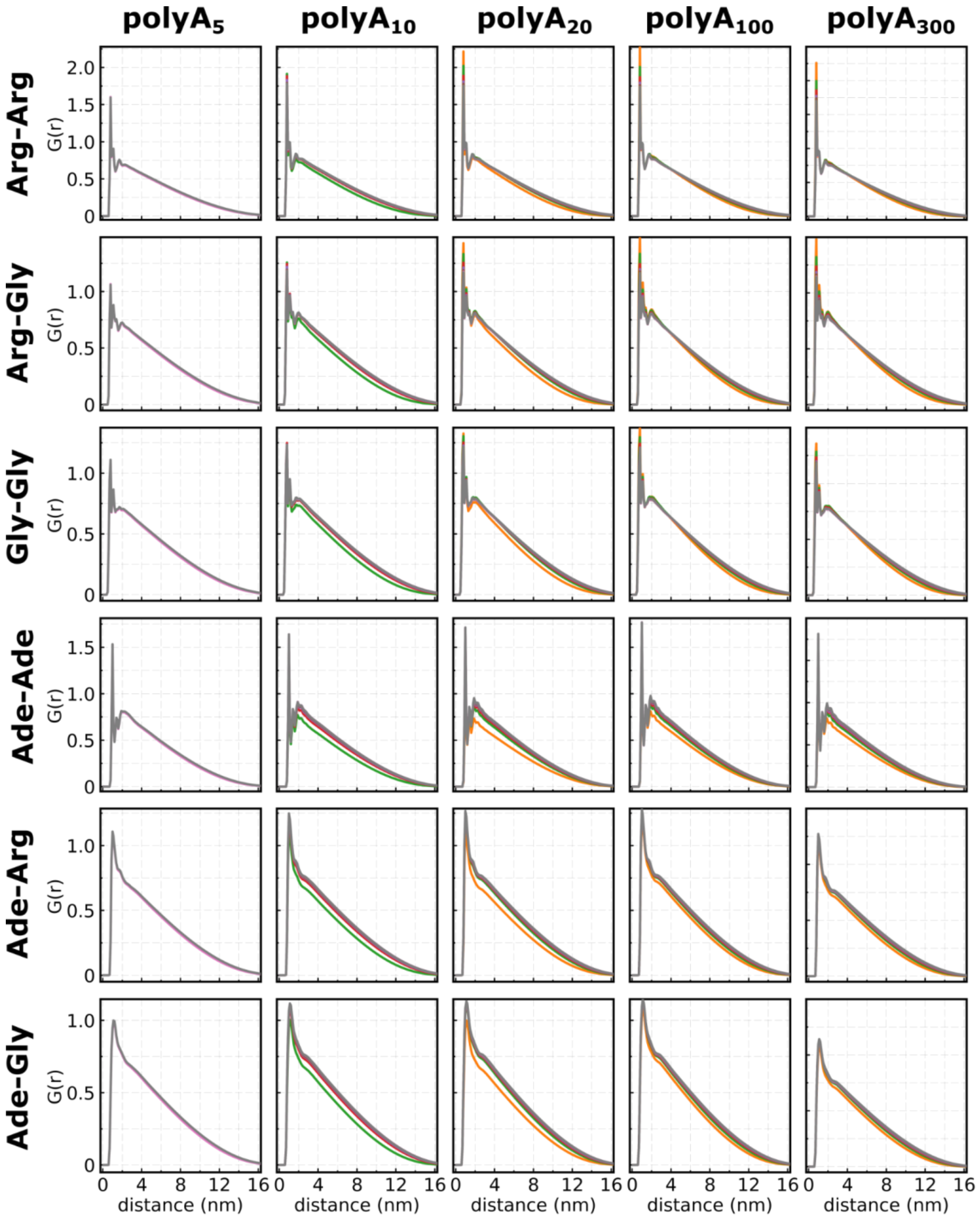


**Figure S4.** Pairwise radial distribution function between different residue types in the condensed phase. Results are shown column-wise for polyA$_N$ (N = 5, 10, 20, 100, and 300) and row-wise for residue pairs (Arg-Arg, Arg-Gly, Gly-Gly, Ade-Ade, Ade-Arg, and Ade-Gly). Each trace shows a different [RGRGG]$_M$ peptide, where M = 1, 2, 3, 4, 5, 6, 8, and 10 are in blue, orange, green, red, purple, brown, and pink, respectively. Initial concentrations for these simulations were kept at 1 mg/mL for both protein and RNA.

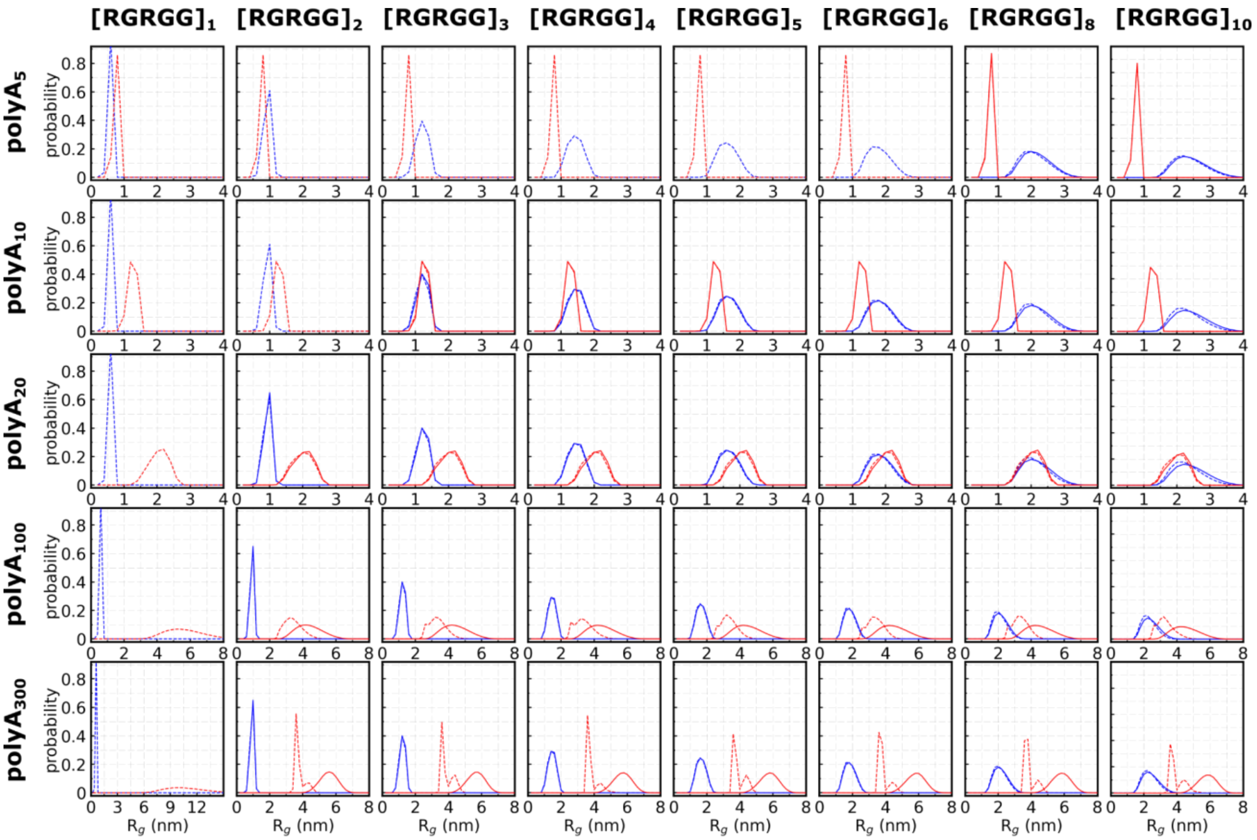


**Figure S5. Distribution of the radii of gyration**. Histograms of radius of gyration ($R_g$) for protein (blue)and RNA (red) when in the condensed (solid line) or disperse (dashed line) phase. Results are shown row-wise for polyA$_N$ (N = 5, 10, 20, 100, and 300) and column-wise for [RGRGG]$_M$ (M = 1, 2, 3, 4, 5, 6, 8, and 10).

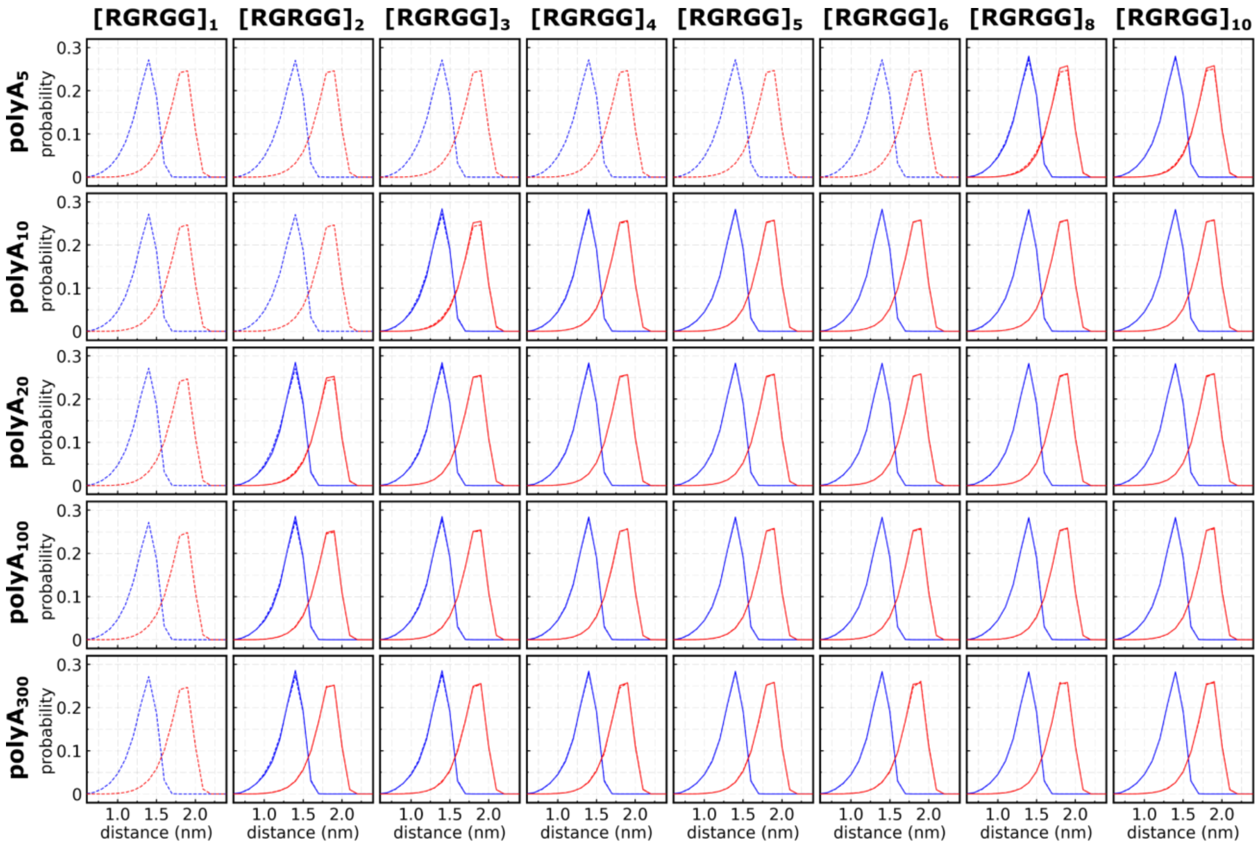


**Figure S6.** Residue 1 to 5 distance distribution. Histograms of distances for Arg1-Gly5 in protein (blue) and Ade1-Ade5 in RNA (red) when in the condensed (solid line) or disperse (dashed line) phase. Results are shown row-wise for polyA$_N$ (N = 5, 10, 20, 100, and 300) and column-wise for [RGRGG]$_M$ (M = 1, 2, 3, 4, 5, 6, 8, and 10).

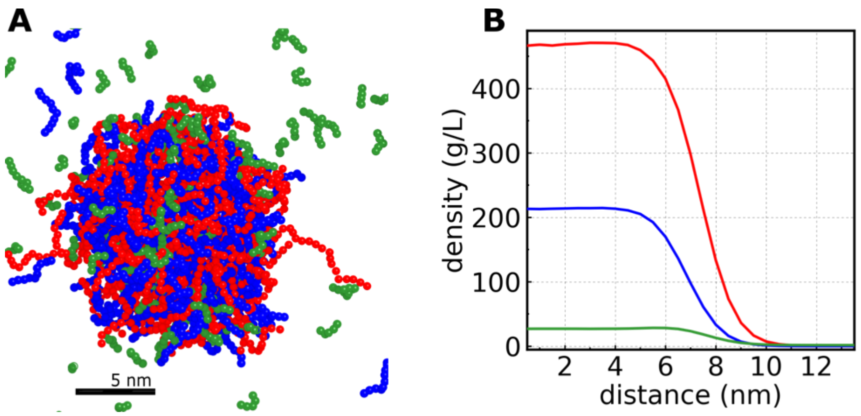


**Figure S7.** Condensate formed in a CG simulation with three components. Both $[RGRGG]_2$ (blue) and $[RGRGG]_1$ (green) are forming the condensate with poly$A_{20}$ (red) (A). The amount of each component inside the condensate is higher than in the dilute phase, as can be seen by the density *vs.* distance from the center of the condensate plot (B). Initial concentrations of poly$A_{20}$, $[RGRGG]_2$, and $[RGRGG]_1$ in this simulation were 1.0, 0.55, and 1.8 mg/mL, respectively. This figure relates to Fig 4 in main text.

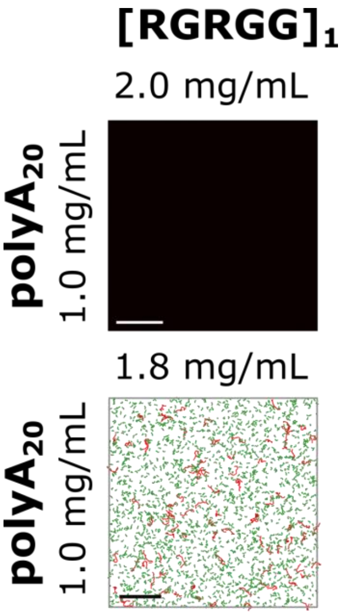


**Figure S8.** LLPS is not observed at higher concentrations of $[RGRGG]_1$. Experimental (upper panel) and simulation (lower panel) results show that $[RGRGG]_1$ alone is not enough to induce phase separation. In the simulation panel, RNA and protein are colored in red and green, respectively. Scale bar represents 10 μm in the experimental panel and 20 nm in the simulation panel. This figure relates to Fig 4 in the main text.
